# Is Kambô psychoactive? Acute and subacute effects of the secretion of the Giant Maki Frog (Phyllomedusa bicolor) on human consciousness

**DOI:** 10.1101/2020.07.22.208223

**Authors:** Timo Torsten Schmidt, Simon Reiche, Caroline L. C. Hage, Felix Bermpohl, Tomislav Majić

**Affiliations:** Department of Psychiatry and Psychotherapy, Berlin Institute of Health, Charité Universitätsmedizin Berlin, corporate member of Freie Universität Berlin, Humboldt-Universität zu Berlin, Campus Charité Mitte, Berlin, Germany; Department of Education and Psychology, Neurocomputation and Neuroimaging Unit (NNU), Freie Universität Berlin, Germany

**Keywords:** Kambô, sapo, Amazonas, Phyllomedusa bicolor, alternative medicine, motivation, Rapé, ayahuasca, Daime, afterglow, psychedelic, Giant Leaf Frog, dermorphin, caerulein, sauvagin, deltorphin

## Abstract

Kambô is the name for the secretion of the Giant Leaf Frog (Phyllomedusa bicolor) containing a plethora of bioactive peptides. Originally, it is ritually used by different ethnicities from the Amazon basin as a remedy against bad luck in hunting. In the last twenty years, Kambô has spread to Western urban centers, often associated with the use of ayahuasca. Anecdotal reports claim beneficial effects on wellbeing and different medical and mental health conditions. However, to date it has been controversial if Kambô elicits altered states of consciousness. Here we retrospectively investigated acute and subacute psychological effects of Kambô in a sample of n = 22 anonymous users (n = 22, mean age: 39 years, ± 8.5; 45.5% female), administering standardized questionnaires for the assessment of psychoactive effects. Acutely, participants reported psychological effects which remained on a mild to moderate level, but no psychedelic-type distortions of perception or thinking. In contrast, persisting effects were predominantly described as positive and pleasant, revealing surprisingly high measures of personal and spiritual significance. Subacute and long-term effects showed some overlap with the “afterglow” phenomena that follow the use of serotonergic psychedelics.

## INTRODUCTION

Kambô is the Matsé name for the secretion of the Giant Leaf Frog (*Phyllomedusa bicolor*), which is ritually used by different ethnicities in the Amazon basin of Brazil and Peru^1^. A variety of potent bioactive peptides have been identified in the frog’s secretion, including phyllocaeruelin, phyllokinin, phyllomedusin, sauvagine, deltorphin^2^, adrenoregulin, and the potent opioids dermorphine and caeruelin^1^. Kambô is obtained from the frog by carefully tying it up and rubbing its skin with a hard instrument, collecting the secretion on a wooden stick. It has been emphasized that in most cases the frog is treated with utmost respect and caution, in order to not harm it, and released it to its natural habitat once that the secretion is collected^3^.

Given its low oral bioavailability, Kambô is most commonly applied by the applicator to the recipient via several fresh superficial burns (“dots”) on the arms, legs or chest^4^. Anecdotally, it has been described that within minutes, the secretion likely enters the lymphatic system and subsequently the blood, thereby inducing an intense reaction that includes hypotension, sweating, tachycardia, heavy vomiting and edema, usually subsiding within an hour. This is followed by listlessness or sleep and, subsequently, a state “perhaps to be described as euphoric”, characterized by increased stamina and clarity of thoughts with an increased capacity for hunting^1^. In Amazonian ethnicities, Kambô is used as a cleansing ritual to liberate hunters from “bad principles” or bad luck in hunting (“panema”), enhancing the recipient’s capacities once that cleansing has occurred and acute effects have subsided^5^.

During the last 20 years, Kambô has found its way to Western urban centers in Brazil and all over the world^6^. Notably, to date none of the substances that have been identified in the Kambô secretion display any serotonergic activity. However, from its Amazonian origins to its use in the context of Brazilian syncretic religions like the Santo Daime and the União do Vegetal^5^ and, finally, to its use in Western healing circles, Kambô has often been associated with the spread of the serotonergic psychedelic ayahuasca^7^. Ayahuasca is an Amazonian shamanic concoction of different plants, including plants (e.g. *Psychotria viridis)* which contain the serotonergic psychedelic N,N-dimethyltryptamine (N,N-DMT) and Banisteriopsis caapi, which contain inhibitors of monoaminoxidase (MAO-I) that render N,N-DMT orally active^8^. Notably, Kambô does not necessarily have to be applied by a shaman and is not considered as a shamanic ritual itself, in contrast to ayahuasca and other ritual plants, where use is restricted to a shamanic framework^9^. During its spread to Western urban centers, however, the Kambô ritual has been transformed from a hunting ritual into therapeutic approaches and a neo-shamanic healing ritual, a process which has been labeled as “shamanization of Kambô”^10^.

The association with ayahuasca, however, is not the only connection between Kambô and nature-derived serotonergic psychedelics. Notably, different names used for the frog’s secretion include “Kambô”, “kampu”, “vaccino da floresta” and also “sapo”, which incorrectly means “toad” in Spanish. This variability of the terms has sometimes led to a confusion of Kambô with the secretion of the Sonoran Desert Toad (Bufo alvarius), which is also referred to as “sapo”^7^. In contrast to Kambô, however, the toad’s secretion contains the potent serotonergic psychedelics 5-methoxy-N,N-dimethyltryptamine (5-MeO-DMT)^11^ and bufotenine, which is usually smoked or snorted, immediately inducing strong psychedelic experiences^12^. Given the different application routes, the two substances are usually not confused by users, even though ceremonies where secretions from Kambô and Bufo alvarius are combined have recently been proposed in Western psychedelic circles.

Another interesting overlap between Kambô and the use of plant-derived psychedelics can be found in anecdotal reports describing beneficial after-effects on wellbeing, medical and mental health problems and personal and spiritual development^13^ – attributes which have previously been associated with the use of serotonergic psychedelics. Of note, the effects of serotonergic psychedelics underlie unique temporal dynamics, with distinct acute (“***psychedelic experiences***” or “***states***”) and subacute effects (“***afterglow***” ***phenomena***)^14^. Afterglow phenomena have been conceptualized as states of “elevated and energetic mood with a relative freedom from concerns of the past and from guilt and anxiety”, which are associated with an enhanced willingness “to enter into close interpersonal relationships”, lasting between two weeks and a month^15^. If these effects are comparable to the aftereffects of Kambô is an open question.

Despite the close cultural and sub-cultural associations between the use of Kambô and different nature-derived psychedelics, no systematic characterization of the acute or subacute effects of Kambô has been reported. Here, we report results of a paper-pencil study among Kambô users employing standardized and validated questionnaires to retrospectively report acute and subacute effects of Kambô. This assessment allows a direct comparison to data from other psychoactive substances and answers in how far the effects of Kambô display similarities with serotonergic psychedelics^16^.

Our study was designed to (1) systematically characterize the acute effects of Kambô, enabling a direct comparison to acute effects of e.g. plant-derived serotonergic psychedelics, (2) explore if Kambô displays subacute effects which might be comparable to the psychedelic afterglow phenomena, including retrospective appraisal of the experiences by the recipients.

## RESULTS

### Sample characteristics

N = 27 datasets were sent back to us, of which n = 5 were excluded due to an insufficient reliability index in the PCI (cut-off h > 2), leaving a final dataset of n = 22 for the first part of the study. The consecutive part of the study on the subacute effects of Kambô was completed by n = 14, where one PEQ dataset and one CS dataset were excluded due to inappropriate completion, leaving for each n=13 datasets.

The sample characteristics are summarized in **Table 1**. With regards to lifetime drug consumption, 15 participants (68.2%) reported experiences with serotonergic hallucinogens (e.g., LSD, DMT, DOM), and 19 (86.4%) reported experiences with ritual plants or traditional indigenous medicines (e.g., ayahuasca, peyote, San Pedro, psilocybin mushrooms, ibogaine, 5-MeO-DMT, bufotenin). Participants reported consumption of other psychotropic substances in their lifetime as follows: cannabis (n = 21, 95.5%), opioids (n = 7, 32.8%), cocaine (n = 10, 45.5%), amphetamine (n = 13, 59.1%), MDMA (n = 15, 68.2%), tranquilizer (n = 4, 18.2%).

**Table 1:**
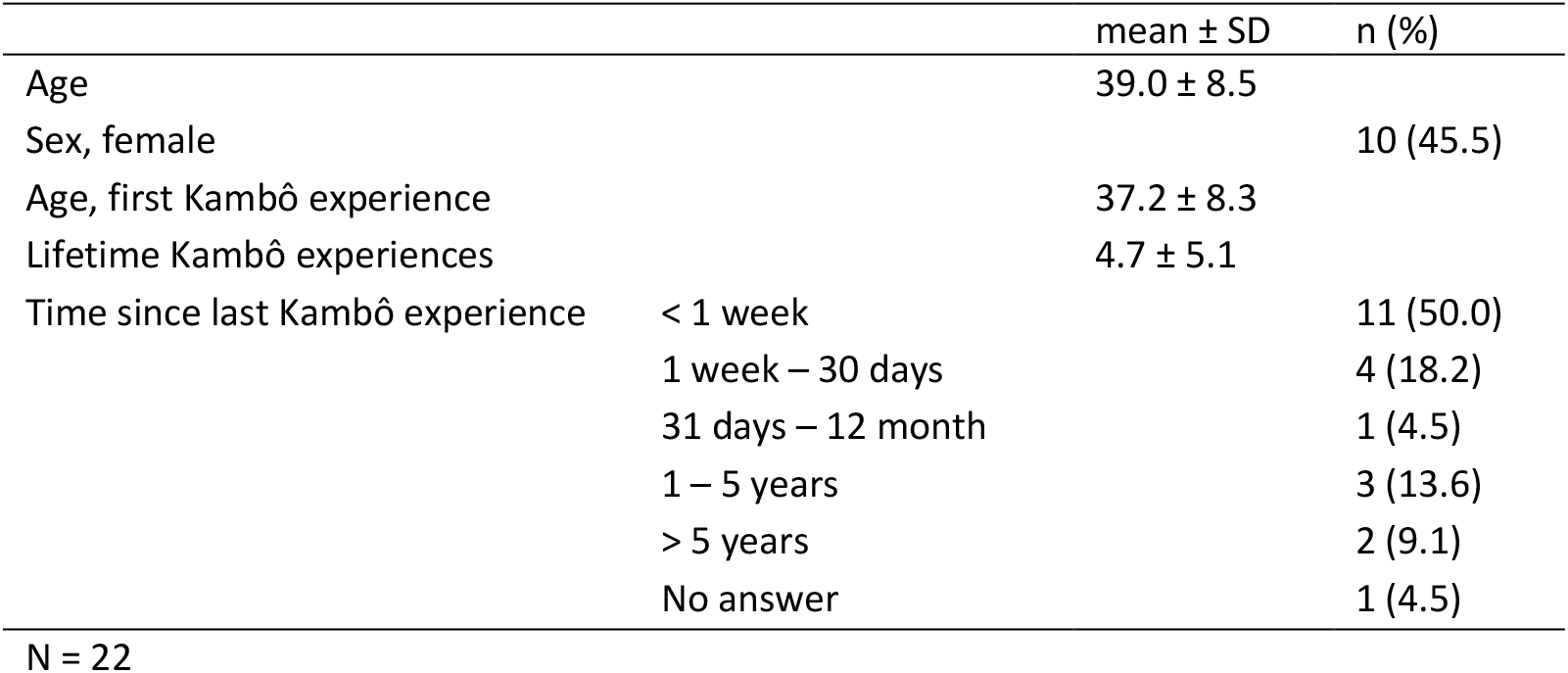
Sample characteristics.

Participants were asked to report on the importance of spiritual practices in their life on a scale from (0 = “not at all” to 100 = “extraordinarily important”), which resulted in an average of 72.7% (± 21.8%).

### Acute subjective effects of Kambô

The Kambô secretion was administered for n = 15 (68.2%) participants by an alternative health practitioner, western healer or “neo-shaman”. In n = 2 (9.1%) cases an indigenous shaman and in n = 3 (13.6%) cases a layperson was reported to have administered Kambô (n = 1 “other” and n = 1 not specified). N = 14 (63.6%) of the participants reported being the only client while receiving Kambô, while n = 7 (31.8%) received it in a group setting (n = 1 not specified). The setting and environment was described as follows: a healing place or temple (n = 11, 52.4%), at home (n = 4, 19.0%), in nature (n = 2, 9.5%), in a ceremony (n = 2, 9.5%), other (n = 2, 9.5%), in a friend’s home (n = 1, 4.8%), with an alternative health practitioner practice (n = 1, 4.8%), at a festival (n = 1, 4.8%), on vacation (n = 1, 4.8%).

Participants reported that Kambô was applied via an average of 7 ± 2 burning points on the participants’ skin. The acute experience lasted less than 15 minutes for n = 6 (27.3%) participants, between 30 and 59 minutes for n = 11 (50.0%) participants, 60 minutes or longer for n = 3 (13.6%) participants (n = 2 not specified). We asked participants explicitly if they thought that Kambô induces a “Rauschzustand” (German word for “inebriation” or “high”), and n = 5 (22.7 %) reported “Yes”.

Acute effects were assessed retrospectively using the ASC rating scale, the PCI, the MEQ and CEQ to assess altered state experiences allowing for direct comparison to hallucinogenic substances and non-pharmacological methods that elucidate psychotropic effects. **Figure 1** and **Table 2** summarize results of the ASC rating scale, the PCI, MEQ30 and CEQ.

**Figure 1:**
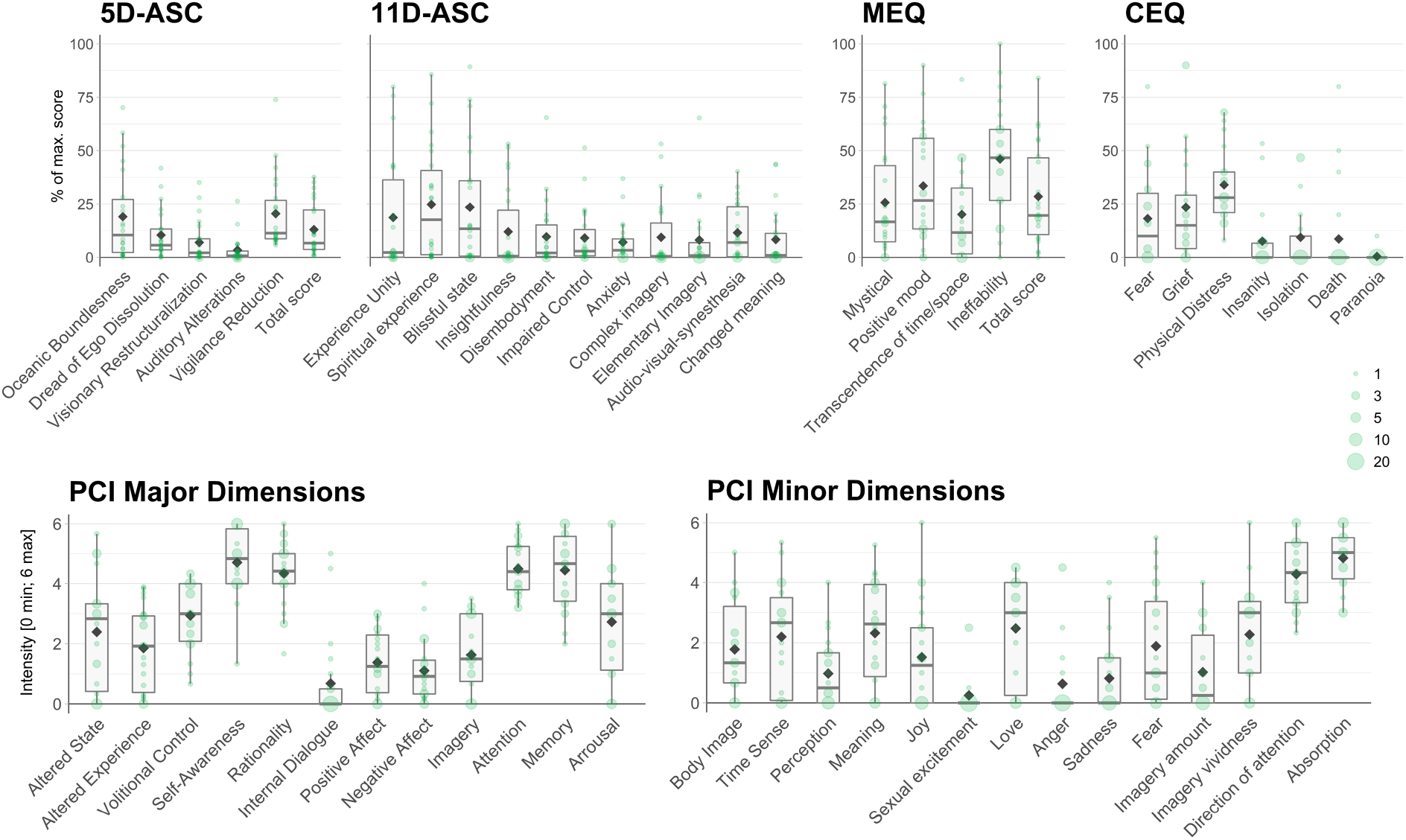
Acute subjective effects of Kambô. Acute effects (n = 22) were assessed with standardized psychometric tools that allow direct comparisons to consciousness alterations induced by different methods as e.g. found on the Altered States Database^16^. Note: The scores of the ASC, MEQ and CEQ are designed to assess differences from zero, while the PCI items are anchored with two opposing statements (See Methods) to the end of the scale [0: minimum; 6: maximum], thereby not assessing differences from zero but characterizing the overall pattern of experience. Note: In comparison to Table 2, the box plots provided here show the data distribution.

**Table 2:** Group level summary statistics for acute effects.

|  | Mean | SD | t-value | p-value |
| --- | --- | --- | --- | --- |
| **5D-ASC** |  |  |  |  |
| Total ASC score | 13.0 | 12.4 | 4.94 | 0.000 |
| Oceanic Boundlessness | 19.1 | 21.2 | 4.22 | 0.000 |
| Dread of Ego Dissolution | 10.5 | 11.5 | 4.26 | 0.000 |
| Visionary Reconstructualization | 7.0 | 10.0 | 3.26 | 0.002 |
| Auditory Alterations | 3.4 | 6.3 | 2.57 | 0.009 |
| Vigilance Reduction | 20.5 | 17.3 | 2.84 | 0.005 |
| **11D-ASC** |  |  |  |  |
| Experience of Unity | 18.7 | 26.8 | 3.27 | 0.002 |
| Spiritual Experience | 24.8 | 26.3 | 4.42 | 0.000 |
| Blissful State | 23.5 | 28.0 | 3.94 | 0.000 |
| Insightfulness | 12.0 | 19.2 | 2.93 | 0.004 |
| Disembodiment | 9.7 | 15.7 | 2.89 | 0.004 |
| Impaired Control and Cognition | 9.1 | 13.6 | 3.15 | 0.002 |
| Anxiety | 7.1 | 9.6 | 3.47 | 0.001 |
| Complex Imagery | 9.4 | 16.4 | 2.68 | 0.007 |
| Elementary Imagery | 8.1 | 15.5 | 2.46 | 0.011 |
| Audio-Visual Synesthesia | 11.6 | 13.1 | 4.15 | 0.000 |
| Changed Meaning of Percepts | 8.3 | 13.9 | 2.82 | 0.005 |
| **PCI** |  |  |  |  |
| Altered State of Awareness | 2.4 | 1.9 |  |  |
| Altered Experience | 1.9 | 1.4 |  |  |
| Volitional Control | 2.9 | 1.1 |  |  |
| Self-Awareness | 4.7 | 1.1 |  |  |
| Rationality | 4.3 | 1.1 |  |  |
| Internal Dialogue | 0.7 | 1.4 |  |  |
| Positive Affect | 1.4 | 1.1 |  |  |
| Negative Affect | 1.1 | 1.1 |  |  |
| Imagery | 1.6 | 1.2 |  |  |
| Attention | 4.5 | 0.9 |  |  |
| Memory | 4.4 | 1.3 |  |  |
| Arousal | 2.7 | 1.9 |  |  |
| Body Image | 1.8 | 1.5 |  |  |
| Time Sense | 2.2 | 1.8 |  |  |
| Perception | 1.0 | 1.1 |  |  |
| Meaning | 2.3 | 1.7 |  |  |
| Joy | 1.5 | 1.7 |  |  |
| Sexual excitement | 2.5 | 1.8 |  |  |
| Love | 0.6 | 1.4 |  |  |
| Anger | 0.6 | 1.4 |  |  |
| Sadness | 0.8 | 1.2 |  |  |
| Fear | 1.9 | 1.9 |  |  |
| Imagery amount | 1.0 | 1.3 |  |  |
| Imagery vividness | 2.3 | 1.7 |  |  |
| Direction of attention | 4.3 | 1.2 |  |  |
| Absorption | 4.8 | 1.0 |  |  |
| **MEQ** |  |  |  |  |
| Mystical | 25.7 | 25.1 | 4.79 | 0.000 |
| Positive Mood | 33.5 | 26.4 | 5.95 | 0.000 |
| Transcendence of time and space | 20.2 | 21.8 | 4.34 | 0.000 |
| Ineffability | 46.1 | 27.0 | 8.01 | 0.000 |
| Total Score | 28.5 | 23.4 | 5.72 | 0.000 |
| **CEQ** |  |  |  |  |
| Fear | 18.2 | 21.9 | 3.53 | 0.001 |
| Grief | 23.5 | 27.1 | 3.68 | 0.001 |
| Physical Distress | 34.0 | 17.9 | 8.07 | 0.000 |
| Insanity | 7.6 | 14.6 | 2.20 | 0.021 |
| Isolation | 9.4 | 17.3 | 2.30 | 0.017 |
| Death | 8.6 | 21.0 | 1.74 | 0.050 |
| Paranoia | 0.5 | 2.1 | 0.90 | 0.189 |

N = 22; To allow direct comparison with previous datasets mean ± SD for all group level scores of the applied questionnaires are provided. For the 5D-ASC, 11D-ASC, MEQ, CEQ scores, we performed one sample t-tests (df = 21) against zero and provide p-values. Please note that data is reported in its completeness and raw significance values are reported instead of significance thresholding. Note: The scores of the ASC, MEQ and CEQ are designed to assess differences from zero, while the PCI items are anchored with two opposing statements (See Methods) to the end of the scale [0: minimum; 6: maximum], therefore not being tested against zero.

### Subacute subjective effects of Kambô

The PEQ and the CS were filled in only when the exemplary Kambô session happened 2 - 3 weeks before filling in the questionnaires. Together these questionnaires cover a broad spectrum of subjectively experienced subacute effects. Results are summarized in **Figure 2** and **Table 3**.

**Figure 2:**
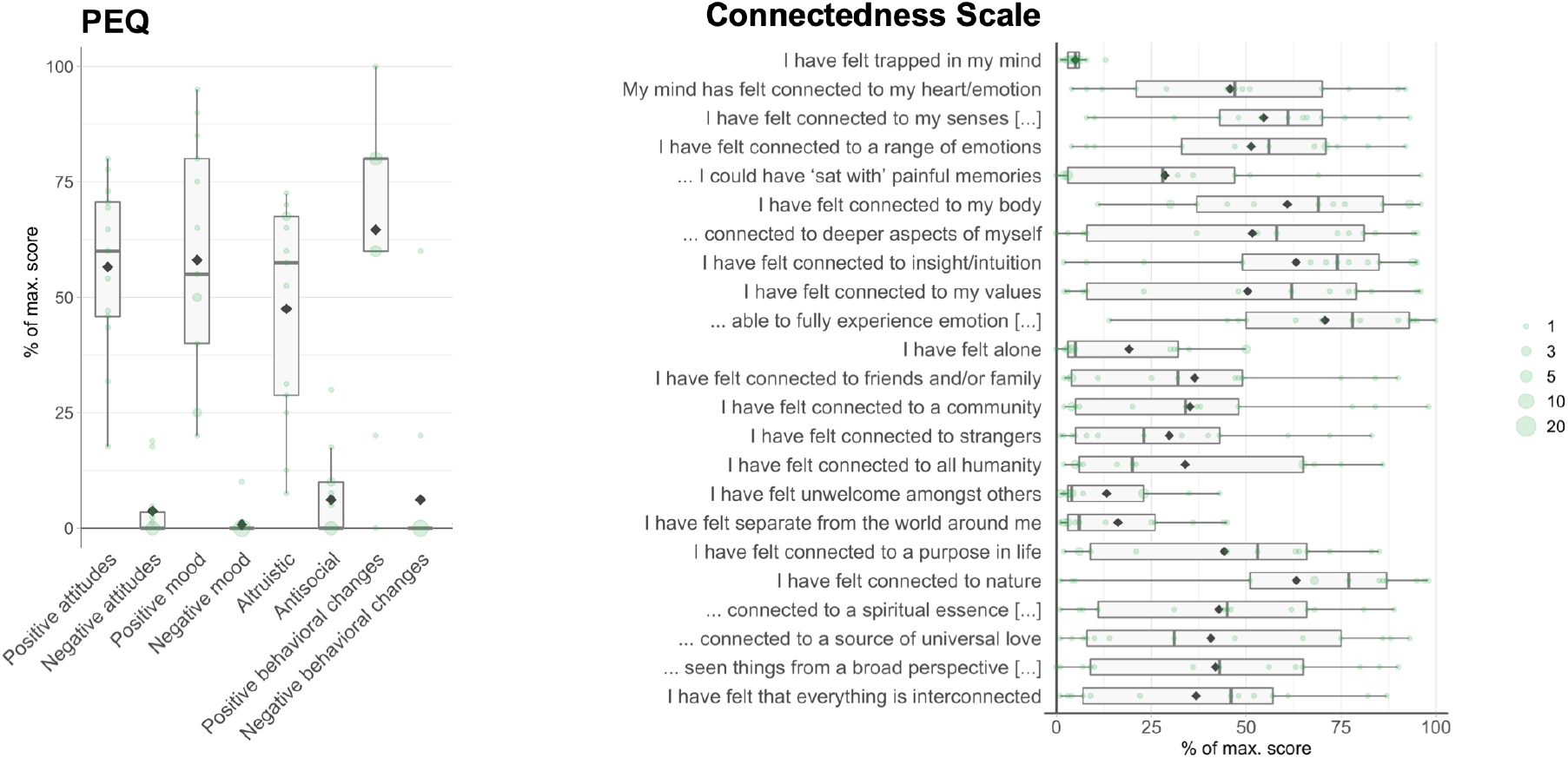
Subacute subjective effects of Kambô. Subacute effects (n = 13) were assessed with the PEQ and the CS and are displayed as box plots for the dimensions or items of the two questionnaires. Note: In comparison to Table 3, the box plots provided here illustrate the data distribution.

**Table 3:**
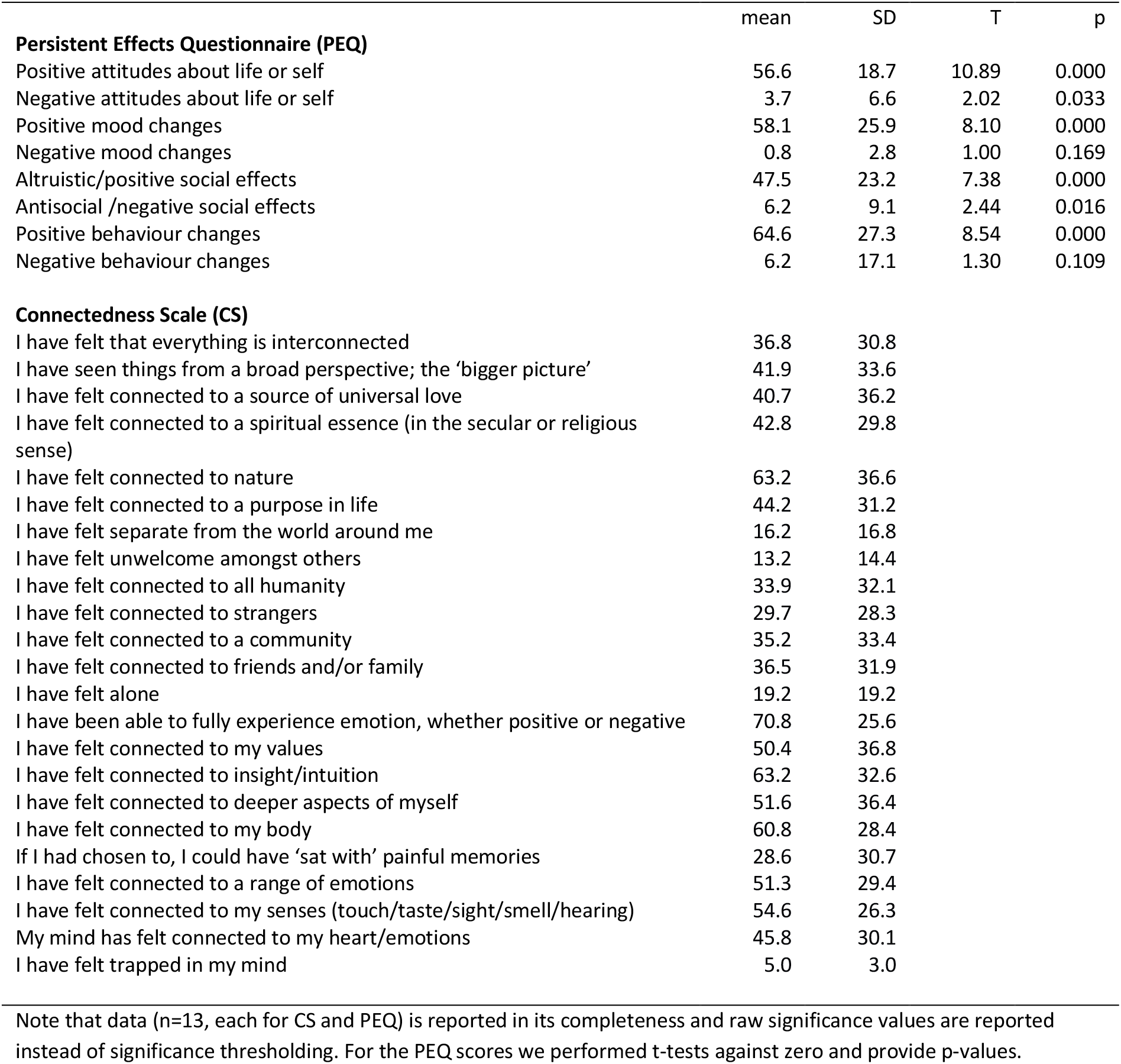
Group level summary statistics for subacute effects.

|  | mean | SD | T | p |
| --- | --- | --- | --- | --- |
| **Persistent Effects Questionnaire (PEQ)** |  |  |  |  |
| Positive attitudes about life or self | 56.6 | 18.7 | 10.89 | 0.000 |
| Negative attitudes about life or self | 3.7 | 6.6 | 2.02 | 0.033 |
| Positive mood changes | 58.1 | 25.9 | 8.10 | 0.000 |
| Negative mood changes | 0.8 | 2.8 | 1.00 | 0.169 |
| Altruistic/positive social effects | 47.5 | 23.2 | 7.38 | 0.000 |
| Antisocial /negative social effects | 6.2 | 9.1 | 2.44 | 0.016 |
| Positive behaviour changes | 64.6 | 27.3 | 8.54 | 0.000 |
| Negative behaviour changes | 6.2 | 17.1 | 1.30 | 0.109 |
| **Connectedness Scale (CS)** |  |  |  |  |
| I have felt that everything is interconnected | 36.8 | 30.8 |  |  |
| I have seen things from a broad perspective; the 'bigger picture' | 41.9 | 33.6 |  |  |
| I have felt connected to a source of universal love | 40.7 | 36.2 |  |  |
| I have felt connected to a spiritual essence (in the secular or religious sense) | 42.8 | 29.8 |  |  |
| I have felt connected to nature | 63.2 | 36.6 |  |  |
| I have felt connected to a purpose in life | 44.2 | 31.2 |  |  |
| I have felt separate from the world around me | 16.2 | 16.8 |  |  |
| I have felt unwelcome amongst others | 13.2 | 14.4 |  |  |
| I have felt connected to all humanity | 33.9 | 32.1 |  |  |
| I have felt connected to strangers | 29.7 | 28.3 |  |  |
| I have felt connected to a community | 35.2 | 33.4 |  |  |
| I have felt connected to friends and/or family | 36.5 | 31.9 |  |  |
| I have felt alone | 19.2 | 19.2 |  |  |
| I have been able to fully experience emotion, whether positive or negative | 70.8 | 25.6 |  |  |
| I have felt connected to my values | 50.4 | 36.8 |  |  |
| I have felt connected to insight/intuition | 63.2 | 32.6 |  |  |
| I have felt connected to deeper aspects of myself | 51.6 | 36.4 |  |  |
| I have felt connected to my body | 60.8 | 28.4 |  |  |
| If I had chosen to, I could have 'sat with' painful memories | 28.6 | 30.7 |  |  |
| I have felt connected to a range of emotions | 51.3 | 29.4 |  |  |
| I have felt connected to my senses (touch/taste/sight/smell/hearing) | 54.6 | 26.3 |  |  |
| My mind has felt connected to my heart/emotions | 45.8 | 30.1 |  |  |
| I have felt trapped in my mind | 5.0 | 3.0 |  |  |

Note that data (n=13, each for CS and PEQ) is reported in its completeness and raw significance values are reported instead of significance thresholding. For the PEQ scores we performed t-tests against zero and provide p-values.

When asked about the spiritual relevance of the Kambô experiences, n = 6 of 13 (46%) participants rated it as strongly spiritually significant, including n = 2 participants who rated it as the single most or among the five most spiritually significant experiences of their life. When asked how personally meaningful the experience was, n = 7 of 13 (54%) participants rated the experience among the ten most meaningful experiences of their life and n = 4 rated it among the five most personally meaningful experience. One participant viewed it as the single most personally meaningful experience in his life. Regarding change of well-being or life satisfaction, n = 8 of 13 (62%) participants stated that the experience increased well-being or life satisfaction slightly, and n = 5 of 13 (38%) participants reported an increase in well-being or life satisfaction between ‘moderate’ and ‘very much’.

## DISCUSSION

In our study investigating effects of Kambô – the secretion of the Giant Maki Frog (Phyllomedusa bicolor) – on human consciousness, we report three major findings: (1) Regarding the acute period of the pharmacological action, only subtle effects on consciousness were recorded, not comparable to serotonergic psychedelics with regard to qualitative aspects and intensity; (2) Participants reported positive subacute after-effects, which in some aspects showed overlaps with psychedelic “afterglow” phenomena; (3) About half of the participants retrospectively appraised their experiences with Kambô as both highly spiritually and personally relevant for their lives, analogous to appraisals of mystical experiences reported under high dosages of serotonergic psychedelics.

### The acute effects of Kambô

In the first part of our study, we set out to test if Kambô induces acute altered states of consciousness (ASCs) comparable to psychedelics. The core of psychedelic experience is currently best assessed with the 11D-ASC questionnaire, the most commonly used tool to assess drug-induced ASCs. The 11D-ASC allows for characterization of subjective effects along 11 dimensions of change in conscious experience. Psychedelic experiences are typically characterized by high scores on the scales *Elementary* and *Complex Imagery, Insightfulness, Spiritual Experience, Experience of Unity* and *Blissful State*^16^. In contrast, for Kambô we found only relatively low scores on these dimensions. Thus, our study demonstrates with standardized questionnaire data that Kambô does not elicit effects on consciousness comparable to serotonergic psychedelics, especially no comparable effects are reported on perception and thinking.

Moreover, we applied the Mystical Experience Questionnaire (MEQ). This tool is used to test for the occurrence of spiritual/mystical experiences, which are thought to have the potential to facilitate conversions, attitude-changes, or even life-changes under special circumstances. High values of “full-blown” mystical experiences have been reported for several serotonergic psychedelics including LSD^17^, psilocybin^18^, and 5-methoxy-dimethyltryptamine^19^. In our sample, few participants reported pronounced aspects of mystical experiences, reflected by relatively low mean scores on the four MEQ factors, with a few higher outlier datapoints. Even if Kambô effects cannot be compared to intense mystical experiences as reported after the use of serotonergic psychedelics^20^ with regard to intensity and completeness of the mystical state, our finding suggests that participants might have experienced psychological and spiritual effects beyond merely somatic reactions. However, this conclusion remains somewhat speculative, and setting as well as expectational and setting factors could have contributed to this effect.

The Challenging Experience Questionnaire (CEQ) includes self-report items designed for the investigation of challenging experiences under psilocybin and other serotonergic psychedelics. These include fear, grief, physical distress, insanity, isolation, fear of death and paranoia, which are symptoms that can occur in challenging experiences (also referred to as “bad trips”) under serotonergic psychedelics^21^. The challenging experiences reported for acute effects of Kambo were mostly limited to “physical distress”. Challenging experiences of a rather psychological nature were barely reported - suggesting unspecific fearful reactions to the strong vegetative effects, but without induction of psychedelic-type psychological crises including insanity, isolation, death or paranoid ideation in the sense of “bad trips”, reflecting distortions of ego functions and self-processing.

Ratings on the Phenomenology of Consciousness Inventory (PCI) allow for comparison of Kambo experiences with hypnosis or meditation techniques to investigate potential shared aspects. Similar to hypnotic or meditative states, participants reported a reduction of positive and negative affect in their Kambô experiences (Pekala 2017). With regard to the question if or how Kambô elicits psychoactive effects, it is noteworthy that the obtained scores on “self-awareness”, “rationality”, “attention” and “memory” indicate that the participants did not feel confused or muddled, which would be expected from centrally active drugs like alcohol or barbiturates.

Taken together, our study provides standardized data that allows a direct comparison of Kambô experiences to the effects of well-known psychoactive substances. On the physiological level, the acute Kambô experience is dominated by an intensive physical reaction, which is reflected in the reports of “physical distress” by our participants and is also likely to have triggered psychological distress in some participants. The process of characterizing acute Kambô effects with standardized questionnaires in the present study revealed that Kambo induces a state of self-centered inwardness. This state does not have typical characteristics of psychedelic-induced states. Although the pharmacodynamics of the Kambô secretion have only been partially investigated, it has been suggested that Kambô’s pharmacological effects are restricted to the cardiovascular, gastroenterological, endocrine and immune systems, the autonomic nervous system (ANS) and the endogenous opioid system^9^. On the one hand, this appears to be plausible given the compounds’ peptide structures which prevent them from passing the blood-brain barrier. On the other hand, the authors hypothesize that the neuropeptide opioids in Kambô (dermorphin, caerulein and deltorphin) could be responsible for the observed “alterations of consciousness”, suggesting psychoactive effects. However, given the neuropeptide structure, these opioids have been reported to be centrally active via intrathecal application only. In addition, the described acute and subacute effects of Kambô resemble stimulant effects rather than those of substances with mu-receptor activity. Thus, the observed acute and subacute effects in our sample are divergent from known psychoactive effects of mu-receptor agonists, suggesting that other compounds or complex interactions between vegetative, neuro-endocrinological and psychological effects might be considered as underlying biological correlates of the Kambô experience. Nevertheless, to date no compounds have been identified that could explain the induction of an ASC during the acute period of Kambô effects and no such phenomena were reported by our participants.

### The subacute effects of Kambô

In the second part of our study, we investigated subacute effects of Kambô up to 2-3 weeks after the reported exemplary session to find first indications if these effects were comparable to psychedelic afterglow phenomena (See^14^). Since no systematic characterization of afterglow phenomena exists to date, even for psychedelics, a quantitative comparison was not possible. Therefore, as a first description of the subacute effects we used the PEQ, previously applied to characterize psychedelic effects mainly in therapeutic contexts. Additionally, we used a questionnaire measuring connectedness, which has not yet been validated.

Notably, subacute effects of Kambô were appraised very positively, including factors of “positive attitudes”, “positive mood”, “altruistic” and “positive behavioural changes”, whereas negative aspects have been reported to be negligible. This is in line with observations describing a euphoric state after the acute effects of Kambô have subsided^1^. Moreover, subacute effects involved dimensions of mood, overall wellbeing, but also aspects on a spiritual and transpersonal level. The intensity of subacute positive effects was pronounced, even if not as intense as after-effects of lysergic acid diethylamide (LSD)^22^ or psilocybin^23^.

Interestingly, the ratings of items describing connectedness to *internal* aspects of oneself were high in our sample, such as being connected to “my senses”, “a range of emotions”, “my body”, “deeper aspects of myself” and “insight/intuition”, and to “have been fully able to experience emotion, whether positive or negative”. In contrast, ratings of items referring to connectedness to *external* aspects (e.g. “a community”, “strangers”, “all humanity”, “a purpose in life”, “spiritual essence” and “a source of universal love”) were far less pronounced, except for the experience of being “connected to nature”. This is in line with anecdotal observations including participants’ subjective experiences of an active interaction with a frog’s “spirit”, which detects and eliminates toxins and bad energy from their mind and body^9^. However, this finding is only partially comparable to mystical experiences associated with acute and subacute effects of serotonergic psychedelics, where states of increased connectedness to both the self and other beings have been reported^23^(i.e. the notion that “everything is interconnected”).

Notably, even if the subacute effects were not comparable to those reported after the use of serotonergic psychedelics regarding intensity and qualitative aspects^22,23^, some of the phenomena which outlasted the acute effects were surprisingly intensive and complex, showing overlaps with psychedelic “afterglow phenomena”, including increases in positive mood, behavior, attitudes and social interaction.

### Limitations

The effects of Kambô reported by our study participants could partially be related to the ritualistic setting of consumption. Our data were collected retrospectively from a group of Western Kambô users recruited through a public workshop on Kambô and a group of practitioners devoted to a specific ritual setting. This might have induced a bias of expectations or motivations for use and thereby involved the placebo dimension. The observed variability in the assessed acute effects suggests that expectational factors and setting might have played a role for some reports. In order to make final conclusions about the psychoactive properties of Kambô, randomized placebo-controlled trials are necessary.

### Conclusion

Our findings demonstrate that the acute effects of Kambô are very different from the effects of serotonergic psychedelics. While the acute effects of Kambô are dominated by strong physical reactions followed by a state of increased inwardness, psychedelic effects appear to facilitate loosening of ego barriers and increased connectedness with oneself and the outer world. Our findings are congruent with anecdotal reports that the subacute effects of Kambô include feelings of being energized with increased stamina and clarity of thoughts, following an initial state of physical sickness and exhaustion. Nevertheless, subacutely, Kambô does exhibit some overlap with serotonergic psychedelics in regard to the reported “afterglow” phenomenon^15^. This finding is striking given the unique temporal dynamics of subacute psychedelic effects, incomparable to any other group of psychoactive substances. Kambô thereby appears to be associated with afterglow-like effects, but without preceding psychedelic states. In agreement with our findings, it has been suggested that the transformative and transpersonal effects of Kambô might be comparable to those associated with the use of serotonergic psychedelics^9^.

## METHODS

### Participants and Procedure

All data of this study was collected anonymously. Potential participants were recruited at a drug information event in Berlin and through Kambô practitioners who forwarded study material to their clients. Participants were informed about the study aim and that data collection is fully anonymously. They were handed out a printed set of paper/pencil-questionnaires together with a prepaid envelope for returning completed sets of questionnaires and gave consent by filling the questionnaire and sending it back anonymously. All procedures were conducted in accordance with the Declaration of Helsinki and were approved by the Ethics committee of the Charité Universitätsmedizin Berlin (EA2/185/17). All questionnaires were applied in German. The first set of questions addressed person specific characteristics, such as age, gender and drug consumption history.

Apart from questions referring to demographic information, a set of questionnaires was given to the participants that comprised the following two parts: (1) questionnaires on the acute effects of an exemplary Kambô session, (2) questionnaires on subacute effects of the exemplary Kambô session. All participants were requested to fill in demographic information and (1). Participants were asked to fill in (2) only if the exemplary Kambô session that they reported about in (1) happened between 2 - 3 weeks ago, as the questions on the acute effects addressed this period after the Kambô session.

As exclusion criterion for data of insufficient quality we used the reliability index (h) of the Phenomenology of Consciousness Inventory (PCI), which is a measure for the participants consistency in completing the questionnaire^24^.

### Assessment of acute subjective effects of Kambô

The second set of questions comprised four well established and validated psychometric tools to assess acute subjective experiences of consciousness alterations:

#### Altered States of Consciousness (ASC) Rating Scale

The ASC rating scale^25,26^ originated from two former versions, the initial APZ (Abnormal Mental States; GERMAN: Abnorme Psychische Zustände)^27–29^ and the revised version OAV^30^ and has become one of the most frequently used questionnaires in the assessment of altered states of consciousness phenomena. The ASC rating scale is supposed to investigate characteristics of consciousness alterations that are invariant across various conditions including both pharmacological (e.g., psilocybin, ketamine, DMT) and behavioral induction methods (e.g., sensory deprivation, hypnosis, autogenic training). Over the course of more than 30 years, the questionnaire underwent several refinements finally leading to the currently used version which comprises 94 items^25,26^. Currently two different analysis schemata are used: the original 5-dimensional scheme (5D-ASC) and the 11-dimensional scheme (11D-ASC)^31^.

#### Phenomenology of Consciousness Inventory (PCI)

The PCI was developed in the context of an interdisciplinary approach described as empirical-phenomenology^24,32^. Most notably influenced by C. T. Tart’s (1975) conception of ASCs, where different states are characterized by distinct structures and patterns of the subjective experience^24^ (p. 192). The PCI assesses subjective experiences along multiple dimensions, where corresponding scales were constructed on the basis of several cluster and factor analyses. Items are presented as two opposing statements (e.g. “I felt ecstatic and joyful” – “I felt no feelings of being ecstatic and joyful”) located on the two poles of a 7-point Likert scale. We used the German version by^33^.

#### Mystical Experiences Questionnaire (MEQ)

The MEQ was first used in the ‘Good Friday Experiment’^34,35^, where it was intended to assess differences regarding aspects of mystical experience between a group taking psilocybin and a control group taking a placebo. Since then, the MEQ has been applied as an instrument for the quantitative assessment of pharmacologically induced mystical experience. Items of the MEQ were chosen based on literature about mysticism including first-person accounts as well as theoretical work, most notably by W. James (1902) and W. T. Stace (1960). The initial MEQ has been further developed by Pahnke (1969), Richards (1975), Griffiths et al. (2006; 2011), and MacLean et al. (2012). The most recent version is the MEQ30^36^, a condensed version of the MEQ with thirty items and four empirical scales^37 38^.

#### Challenging Experiences Questionnaire (CEQ)

The CEQ was designed to provide a tool for the comprehensive assessment of acute negative effects of temporarily induced altered states of consciousness^39^, based on three questionnaires: Hallucinogenic Rating Scale (HRS), 5D-ASC, and States of Consciousness Questionnaire (SOCQ). The conceptual scope of the 26-item CEQ is informed by literature on psychological and physical distress following hallucinogen intake and covers a variety of adverse cognitive, affective, and somatic reactions which are clustered into seven distinct dimensions of challenging experiences^37^. Analyzing data from an online survey on negative experiences with psilocybin, corresponding scales for the first six dimension were derived by exploratory factor analysis, complemented by the Paranoia scale, and subsequently validated through confirmatory factor analysis^39^. Items are rated on a 6-point scale adopted from the SOCQ.

### Assessment of subacute subjective effects of Kambô

The third set of questionnaires addressing subacute effects was introduced with the instructions to fill out the following questionnaires only if the exemplary Kambo session (for which the acute effects were reported) had happened up to 3 weeks before the day of completing the questionnaire. The set of questions comprised the following two questionnaires:

#### Items about Connectedness

We used a previously unpublished set of 23 questions to measure different aspects of connectedness^40^. The items include aspects of connectedness to one’s self, connectedness to the universe and connectedness to others. The items stem from the development of a *connectedness scale (CS)*. As no validation or confirmation of a factor structure had been published until now, the scores on this scale are presented for the individual items.

#### Persisting Effects Questionnaire (PEQ)

The PEQ was developed as a follow-up questionnaire to the States of Consciousness Questionnaire (SOCQ), which was an intermediate version of the MEQ^20^. The PEQ assesses long-term changes in participants. We used a German version of the extended PEQ version reported in^23^. This version uses 140 items rated on a 6-point Likert scale from 0 (‘not at all’) to 5 (‘extremely’), and 3 items on the retrospective assessment of the importance and effects of the experiences.

### Data Analysis and Visualization

Questionnaire data were analyzed using standardized analysis sheets. Data visualization was performed with RStudio v.1.2.1335 and package ggplot2 v.3.2.1. Boxplots represent the lower and upper hinges corresponding to the first and third quartiles (the 25th and 75th percentiles), the median as a horizontal bar and the whiskers indicate the range of the data (limited to 1.5 x interquartile range). Discrete individual data points are shown as green dots. To display overlapping data points the size of the dots represents the number of observations at each value. Data points not lying within the range of the whiskers are outliers. The mean, without any outlier exclusion, is displayed as a diamond shape. For comparability with previous reports, the means and SD for all questionnaire data are provided in **Table 2** (without outlier exclusion).

## Acknowledgements

The authors would like to thank Tobias Thon† for help in recruitment of participants and Amy Romanello for proof reading of the manuscript.

## Data Availability

Data is available upon personal request.

## Author Contributions Statement

T.T.S. and T.M. designed the study. C.H., S.R. and T.T.S. analyzed the data, S.R. designed figures and tables and T.T.S., T.M. and F.B. wrote the manuscript. All authors edited the manuscript and approved its final version. All authors agreed to be personally accountable for their own contributions and to ensure that questions related to the accuracy or integrity of any part of the work, even ones in which the author was not personally involved, are appropriately investigated, resolved, and the resolution documented in the literature.

